# Integrated multiomic profiling of *SCN2A* loss-of-function reveals widespread molecular remodeling in patient hiPSC-derived neurons

**DOI:** 10.64898/2026.03.20.713167

**Authors:** Adne Vitória Rocha Lima, Erik Aranha Rossi, Izabela Mamede, Gisele Vieira Rocha, Thaís Alves de Santana, Elisama Araújo da Silva, Rachel Santana Cunha, Fernanda Martins Marim, Victor Emmanuel Viana Geddes, Paola Alejandra Fiorani Celedon, Carolina Kymie Vasques Nonaka, Kátia Nunes da Silva, Dalila Lucíola Zanette, Zaquer Suzana Munhoz Costa-Ferro, Clarissa Araújo Gurgel Rocha, Renato Santana Aguiar, Yang Yang, Bruno Solano de Freitas Souza

**Author notes:** Correspondence (B.S.F.S.). These authors contributed equally and share first authorship: AVRL & EAR.

## Abstract

*SCN2A*-related neurodevelopmental disorders comprise a genetically and mechanistically diverse group of early-onset brain conditions. Loss-of-function (LoF) variants in *SCN2A* represent one of the strongest genetic risk factors for autism spectrum disorder and intellectual disability, yet the molecular cascade linking reduced NaV1.2 dosage to neuronal dysfunction remains poorly understood. Here, we combine deep isoform-resolved transcriptomics, high-content imaging, and high-content cellular phenotyping in human hiPSC-derived neurons from three unrelated individuals carrying pathogenic *SCN2A* LoF variants and three independent healthy donor lines to delineate the multi-layered consequences of NaV1.2 insufficiency. We show that *SCN2A* LoF activates the nonsense-mediated decay (NMD) mechanism, selectively depleting canonical *SCN2A* isoforms and modifying the cell’s RNA processing. These molecular deficits translate into robust structural phenotypes, including axon initial segment shortening, reduced sodium channel density, and simplified dendritic arborization. Transcriptomic analysis converged on remodeling of synaptic and axonal pathways. RNA-seq identified coordinated alterations in gene programs linked to synaptic signaling, ion channel activity, and neuronal projection development, consistent with the structural and functional phenotypes observed. Transcript-level analysis further uncovered extensive perturbation of long non-coding RNA (lncRNA) networks, including lncRNAs strongly correlated with *SYN1* and *ANK3* isoforms. Together, these findings reveal that *SCN2A* haploinsufficiency induces a phenotype spanning NMD activation, isoform-specific dysregulation, axon initial segment destabilization and lncRNA-dependent regulatory shifts. This multiscale framework clarifies how reduced NaV1.2 disrupts neuronal development and highlights isoform-level restoration and modulation of post-transcriptional control as promising therapeutic avenues for *SCN2A*-related neurodevelopmental disorders.

## Introduction

*SCN2A*-related neurodevelopmental disorders (*SCN2A*-NDDs) represent one of the most mechanistically stratified genetic groups within early-onset brain conditions (Sanders et al., 2018; Satterstrom et al., 2020). Pathogenic variants in *SCN2A*, which encodes the voltage-gated sodium channel NaV1.2, give rise to a clinically heterogeneous spectrum ranging from neonatal epileptic encephalopathies to global developmental delay, intellectual disability, and autism spectrum disorder (ASD) (Reynolds; King; Gorman, 2020; Thompson et al., 2023). This phenotypic diversity is driven by variant-specific functional effects: gain-of-function (GoF) mutations typically increase sodium current and lead to early-life seizures, whereas loss-of-function (LoF) variants reduce NaV1.2 activity and are predominantly associated with ASD and cognitive impairment without seizures (Ben-Shalom et al., 2017; Wolff; Brunklaus; Zuberi, 2019). This bidirectional genotype–phenotype mapping has established *SCN2A* as a defining locus in precision neurogenetics and as a key point for understanding the molecular architecture of neurodevelopmental dysfunction.

NaV1.2 plays an essential role in action potential initiation, axon initial segment (AIS) excitability, dendritic backpropagation, and early circuit maturation (Harley et al., 2023; Hu et al., 2009). Its expression peaks during perinatal cortical development, a critical window for establishing excitatory–inhibitory balance and synaptic refinement (Ogiwara et al., 2018; Zhang et al., 2021). Accordingly, perturbations in NaV1.2 dosage disrupt network formation, providing a direct mechanistic link between *SCN2A* haploinsufficiency and neurodevelopmental phenotypes (Spratt et al., 2019; Tamura et al., 2025). Human genetic studies have consistently placed *SCN2A* among the highest-effect genes for ASD, with LoF variants conferring some of the strongest monogenic risks identified to date (De Rubeis et al., 2014; Sanders et al., 2015).

A major subset of pathogenic *SCN2A* LoF variants operates at the RNA level through introduction of premature termination codons (PTCs) that engage the nonsense-mediated mRNA decay (NMD) pathway. NMD is a translation-coupled surveillance system that selectively degrades PTC-containing transcripts, serving as a potent mechanism for reducing gene dosage (Carrard; Lejeune, 2023; Kurosaki; Popp; Maquat, 2019). Recent work in engineered stem cell and mouse systems has validated this mechanism, demonstrating that NMD-driven transcript depletion impairs sodium channel density at the AIS, dampens neuronal firing, and alters synaptic maturation (Asadollahi et al., 2023; Chen et al., 2024; Tamura et al., 2025)).

Despite these advances, key mechanistic questions remain unresolved. In particular, most prior studies have relied on engineered mutations introduced into reference cell lines or animal models, which do not capture the complete genomic, regulatory, and epigenetic context of affected individuals (Avior; Sagi; Benvenisty, 2016; Soldner; Jaenisch, 2018). Human induced pluripotent stem cell (hiPSC)-derived neurons overcome these limitations by preserving the endogenous variant, its native transcriptomic environment, and patient-specific compensatory responses (Sergiu P. Pașca, 2018; Volpato; Webber, 2020). Patient-derived *SCN2A* LoF neuronal models offer a unique platform to dissect how NMD shapes isoform usage, protein abundance, subcellular organization, and neuronal morphology. However, no study to date has integrated isoform-resolved transcriptomics and high-content neuronal phenotyping to define the multi-layer consequences of *SCN2A* haploinsufficiency. Here, we address these gaps by leveraging a set of hiPSC-derived neurons from patients carrying distinct *SCN2A* LoF variants predicted to activate NMD. Using a multi-omic strategy combining isoform-level RNA sequencing, immunocytochemistry, and high-content morphological profiling, we dissect the molecular and cellular architecture of *SCN2A* haploinsufficiency and define a convergent multi-scale phenotype of *SCN2A* LoF in human neurons, revealing coordinated disruptions in synaptic scaffolding, ion channel complexes, dendritic development, and intracellular trafficking. This multi-omic approach complements existing functional studies by defining the upstream molecular and structural alterations that underlie the well-documented electrophysiological deficits in *SCN2A* LoF models.

## Methods

### Study design and ethical approval

This study included six previously characterized hiPSC lines, three unaffected, unrelated controls (CTL-1,CTL-2,CTL-3); (Paredes et al., 2019) and three patient-derived lines carrying *SCN2A* LoF variants, clinically presenting with severe ASD and intellectual disability, without seizures. Two patients carried a heterozygous nonsense variant (p.Arg856Ter; SCN2A-1 and SCN2A-3), and one carried a heterozygous frameshift variant (p.Glu169Aspfs*13; SCN2A-2), as previously characterized by our group (Sampaio et al., 2019; Santos et al., 2025). Variants were classified as LoF based on predicted premature truncation and prior clinical and molecular characterization. All procedures were conducted in accordance with institutional guidelines and were approved by the Research Ethics Committee of Hospital São Rafael (CAAE: 51802521.6.0000.0048).

### HiPSC culture, generation of Ngn2-inducible lines and characterization

The hiPSCs were thawed from cryopreserved stocks prepared in KnockOut Serum Replacement (KSR; Thermo Fisher Scientific) supplemented with 10% dimethyl sulfoxide (DMSO; OriGen). The cells were plated onto Geltrex-coated six-well plates and maintained in StemFlex medium (Thermo Fisher Scientific) at 37 °C and 5% CO₂, with medium changes every 48 h. Cultures were inspected daily for morphology and confluency using an inverted microscope (Nikon Eclipse Ti-U). At 80–90% confluency, the cells were dissociated with Accutase or Versene (Thermo Fisher Scientific) and replated at a 1:5 ratio for expansion or cryopreservation. Thiazovivin (5 µM; Tocris) was added post-passaging to enhance cell survival. Routine mycoplasma screening was performed using a PCR-based assay as described previously (Martins et al., 2019).

Doxycycline-inducible Ngn2 hiPSC lines (hiPSC-Ngn2) were generated by lentiviral transduction as described in (Schafer et al., 2019). Lentiviral particles were produced in HEK293FT cells by co-transfecting pMD2.G (#12259, Addgene), psPAX2 (#12260, Addgene), and pLVX-UbC-rtTA-Ngn2:2A:EGFP (#127288, Addgene) using the calcium phosphate method, as described previously (Souza et al., 2017). Viral supernatants were collected at 48 and 72 h post-transfection, pooled, and concentrated by ultrafiltration (Amicon Ultra, 30 kDa; Millipore). HiPSCs were transduced with concentrated viral particles in the presence of polybrene (6 µL/mL; Sigma Aldrich #TR1003) and selected with puromycin (5 µg/mL) for 48 h to establish stable hiPSC-Ngn2 clones. Characterization of hiPSC-Ngn2 lines included assessments of pluripotency, neuronal induction and chromosomal integrity, as described in the next sections.

### GTG-Band Karyotyping

Chromosomal stability was assessed by karyotyping using G-banded (GTG) metaphase analysis. HiPSCs were treated with Colcemid (0.1 µg/mL; Thermo Fisher Scientific) for 2 h to arrest cells in metaphase, incubated in hypotonic KCl (0.075 M) at 37 °C for 15 min, and fixed in Carnoy’s fixative (methanol:acetic acid, 3:1) a minimum of three fixation steps at least three fixation steps. Slides were stained and analyzed at 400–500 band resolution using a Zeiss Axiostar microscope. A minimum of 20 metaphases per line were evaluated according to the International System for Human Cytogenomic Nomenclature (ISCN 2020).

### Neuronal differentiation and validation

For neuronal induction, hiPSC-Ngn2 lines were cultured until approximately 70% confluency, then treated with doxycycline (2 µg/mL; Sigma-Aldrich, USA). After 24 h of induction, the culture medium was replaced with Neurobasal Plus supplemented with B27 Plus and N2 (Thermo Fisher Scientific) and Laminin (Gibco). The cells were maintained for 15 days, with medium changes every 48 h. All differentiation batches included the six hiPSC lines (three controls and three *SCN2A* LoF lines), which were processed in parallel under identical conditions. For each assay, replicates correspond to independent differentiation experiments performed on different days. Neuronal identity was confirmed by immunostaining for βIII-tubulin (TUJ1), microtubule-associated protein 2 (MAP2), and synapsin I (SYN1). Nuclei were counterstained with DAPI. Fluorescence images were acquired using a Nikon Eclipse Ti-S inverted microscope equipped with 20× and 40× objectives and processed with NIS-Elements software (Nikon Instruments, Tokyo, Japan).

### Gene expression analyses

Pluripotency of parental and genetically-modified lines was confirmed by RT-PCR analysis of NANOG, SOX2 and OCT4. Total RNA was extracted using the RNeasy Mini Kit (Qiagen), and 1.5 µg of RNA was reverse-transcribed with the High-Capacity cDNA Reverse Transcription Kit (Thermo Fisher Scientific). cDNA amplification was performed using Platinum™ Taq DNA Polymerase (Thermo Fisher Scientific) under standard PCR cycling conditions. For characterization of hiPSC-derived neurons, total RNA was isolated using the RNeasy Mini Kit (Qiagen) and reverse-transcribed with the High-Capacity cDNA Reverse Transcription Kit (Applied Biosystems). Quantitative PCR (qPCR) was performed on an ABI Prism™ 7500 Fast Real-Time PCR System using either SYBR™ Green Master Mix (Thermo Fisher Scientific) or TaqMan™ Gene Expression Assay. Relative gene expression was quantified using the 2⁻ΔΔCt method with ACTB or GAPDH as endogenous control.

### Confocal imaging and image quantification

For immunofluorescence, Ngn2-induced neurons were cultured on Geltrex-coated glass coverslips, fixed with 4% paraformaldehyde (Sigma-Aldrich) for 15 min, permeabilized with 0.3% Triton X-100 for 10 min, and blocked with 3% bovine serum albumin in PBS for 1 h at room temperature. Cells were incubated overnight at 4 °C with primary antibodies (listed in Supplementary Table 1), followed by Alexa Fluor–conjugated secondary antibodies (Thermo Fisher Scientific) for 1 h at room temperature. Nuclei were counterstained with DAPI (1 µg/mL), and coverslips were mounted with Fluoromount-G (SouthernBiotech). Images were acquired on a Nikon A1 laser-scanning confocal microscope using 20× or 40× Plan Apo objectives and processed with NIS-Elements AR software (Nikon Instruments).

Neuronal morphology and protein localization were analyzed using a Leica TCS SP8 laser-scanning confocal microscope (Leica Microsystems, Germany) equipped with a 63× or 100x oil-immersion objective. Acquisition parameters (laser power, gain, pinhole, and offset) were kept constant across all experimental groups to ensure quantitative comparability. Image analysis was performed using ImageJ/Fiji (NIH, USA). Fluorescence intensity of ankyrin-G (AIS marker) and pan-Nav (voltage-gated sodium channels) was quantified using manually defined regions of interest (ROIs). AIS length, mean fluorescence intensity, and integrated density were measured for each ROI and normalized to background-subtracted signal intensity. Only NaV1.2-positive puncta colocalizing with ankyrin-G were included in the analyses. Dendritic complexity was assessed by Sholl analysis, performed as previously described (Costa-Ferro et al., 2025), to quantify neuronal arborization as a function of distance from the soma.

### Cell Painting Assay and High-Content Quantification

Cell Painting was performed to extract morphological and subcellular features from hiPSC-derived neurons using a standardized multiplex staining panel. Cells were labeled with Hoechst (nuclei), Concanavalin A–Alexa Fluor 488™ (endoplasmic reticulum and plasma membrane), WGA–Alexa Fluor 555™ (Golgi/plasma membrane), SYTO™ (RNA-rich regions), MitoTracker Deep Red™ (mitochondria), and phalloidin–Alexa Fluor 568™ (F-actin, cytoskeleton). After staining during 30 minutes in the dark, at room temperature, cells were washed twice with PBS 1X, for acquisition with the CellInsight™ CX7 LZR High-Content Screening Platform (Thermo Fisher Scientific), using a 20x objective to capture 81 fields per well. Automated acquisition with fixed exposure parameters across all conditions was performed.

Image quantification and feature extraction were carried out using the Cell Painting bioapplication. An automatic detection of subcellular structures was performed using the software in order to extract compartment-specific metrics (intensity, texture, granularity, and distribution) and to generate normalized feature matrices for downstream analyses comparing *SCN2A* LoF and control neurons. Principal Component Analysis (PCA) was conducted on the normalized feature matrix after averaging single-cell measurements per well. Data were standardized (mean = 0, SD = 1), and principal components were selected using Prism’s parallel analysis with a 95% threshold and 1,000 simulations in automatic mode. PCA plots were generated in GraphPad Prism using the resulting eigenvalues and loadings.

### Cycloheximide Treatment for NMD Inhibition

To assess the sensitivity of *SCN2A* transcripts to NMD, hiPSC-derived neurons at day 15 of differentiation (D15) were treated with 100 µM cycloheximide (CHX) for 6 h, an adapted version of a previously described protocol (Lizarraga et al., 2021). Control wells received an equivalent volume of vehicle (DMSO), after incubation, cells were processed for RNA isolation and qPCR as described in the RNA extraction and quantitative PCR section. Relative changes in *SCN2A* transcript abundance following CHX treatment were used to infer NMD responsiveness.

### RNA Extraction, Library Preparation and Sequencing

Cells were harvested and total cellular RNA was isolated using the RNeasy Mini kit (Qiagen) according to manufacturer’s instructions. RNA quantification was performed using Qubit RNA HS Assay kit and Qubit 3 Fluorometer (Thermo Fisher Scientific). Genomic DNA contamination was avoided by DNase treatment (TURBO DNA-free Kit, Thermo Fisher Scientific) before RNA-Seq and RT-qPCR validation. Only samples with RNA integrity number (RIN) ≥ 9 were used, as verified by RNA 6000 Pico Kit and 2100 Bioanalyzer (Agilent Technologies). Ribosomal RNA was depleted using Ribo-Zero Gold (Illumina) and 200 ng of total RNA for each sample were used for library preparation with TruSeq Stranded Total RNA Library Prep (Illumina) according to manufacturer’s instructions. RNA-seq were performed in 2 × 9 samples using NextSeq 2000 High Output v2 Kit (150 cycles) (Illumina) in a NextSeq 2000 platform (Illumina).

### Computational Pipeline for Gene- and Transcript-Level Differential Expression and Isoform Analysis

For gene-level differential expression analysis (Figure 4), paired-end FASTQ files were preprocessed using Trimmomatic v0.39 (Bolger; Lohse; Usadel, 2014). Quality assessment of both raw and trimmed reads was performed with FastQC, allowing evaluation of GC content, per-base quality scores, residual adapter contamination, and sequence complexity. Filtered reads were aligned to the human reference genome GRCh38 - GCA_000001405.29 using HISAT2 (Kim et al., 2019), and the resulting SAM files were converted and sorted into BAM format using samtools sort. Gene-level quantification was performed with featureCounts (Subread package) using the annotation file Homo_sapiens.GRCh38.106.gtf. To correct for technical effects and unwanted sources of variation, normalization was conducted using the RUVs method implemented in the RUVSeq package (version 0.4.2) (Molania et al., 2023). Subsequently, differential expression analysis was carried out using the dream statistical framework from the variancePartition package (version 1.36.3) (Hoffman; Roussos, 2021), which incorporates variance weights estimated by voomWithDreamWeights and accommodates both fixed and random effects. Differentially expressed genes (DEGs) were defined using the thresholds FDR < 0.05 and |log₂FC| > 0.5.

To contextualize the transcriptional profiles of our hiPSC-derived neurons within human brain development and external reference datasets, we performed a comparative mapping analysis using publicly available transcriptomic compendia BrainSpan brain samples. Gene-level normalized expression matrices from these databases were harmonized to our dataset by matching gene identifiers and filtering for shared, high-confidence genes. Samples from each reference dataset were restricted to cortical regions and developmental stages relevant to early neurogenesis. Multidimensional scaling (MDS) and principal component analysis (PCA) were then computed jointly using variance-stabilized counts from our samples and the external datasets, enabling projection of *SCN2A* LoF and control neurons onto the developmental landscape.

For transcript-level quantification (Figures 2 and 6), samples were pseudo-aligned using Salmon (Patro et al., 2017) against the most recent reference human transcriptome annotation from GENCODE (GENCODE vh49). The output from Salmon was then run through Terminus (Sarkar et al., 2020), which summarizes transcripts that are too similar to be distinguished on each specific sample. The quant files were then imported into R using tximport (Soneson; Love; Robinson, 2015)) and tximeta (Love et al., 2020).

Differentially expressed transcripts (DETs) were calculated using the swish function of package Fishpond (Zhu et al., 2019). Data was processed and visualized in R using packages tidyomics (Hutchison et al., 2024), ggplot2 (Valero-Mora, 2010) and EnhancedVolcano and further isoform visualization was performed using Isoformic (Mamede et al., 2025). Guilt-by-association analysis of lncRNA isoforms and protein coding isoforms were performed using spearman correlation test between lncRNA-protein coding pairs using the TPMs of all the samples. Correlations with FDR lower than 0.01 and absolute rho value higher than 0.8 were marked as significant.

## Results

### NMD underlies reduced *SCN2A* dosage in patient hiPSC-derived neurons

To establish patient-derived neuronal models of *SCN2A* LoF, we first characterized three patient hiPSC lines modified for dox-inducible Ngn2 expression, carrying heterozygous pathogenic variants: two with nonsense mutations (p.Arg856; SCN2A-1 and SCN2A-3) and one with a frameshift mutation (p.Glu169Aspfs13; SCN2A-2). These variants are located in distinct domains of the NaV1.2 channel, the frameshift variant in Domain 1 and the nonsense variants in Domain 2, both predicted to generate PTCs that activate NMD. All hiPSC lines displayed typical colony morphology, expressed key pluripotency markers (OCT3/4, Nanog, Sox2, Tra-1-60), and maintained normal karyotype (Supplementary Figure 1).

The structural schematic (Figure 1A) illustrates the four homologous domains (D1-D4) of the NaV1.2 channel, each containing six transmembrane segments (S1-S6). Both variant types introduce PTCs predicted to engage NMD, resulting in transcript degradation and *SCN2A* haploinsufficiency. Using a dox-inducible Ngn2 system, we differentiated hiPSCs into cortical neurons over a 15-day protocol (Figure 1B). Neuronal identity was confirmed by immunostaining for MAP2, NEUN, and TUJ1 (Figure 1C), which demonstrated proper neuronal morphology in both control and mutant *SCN2A* lines. Expression levels of MAP2 and vGLUT1 mRNA did not differ between groups (Figure 1D-E), indicating equivalent differentiation efficiency. Joint MDS with BrainSpan positioned the generated *SCN2A* and control hiPSC-neurons within the early/mid-fetal cortical cluster, showing that their transcriptomic profiles most closely match mid-fetal stages of human cortical development (Figure 1F).

**Figure 1.**
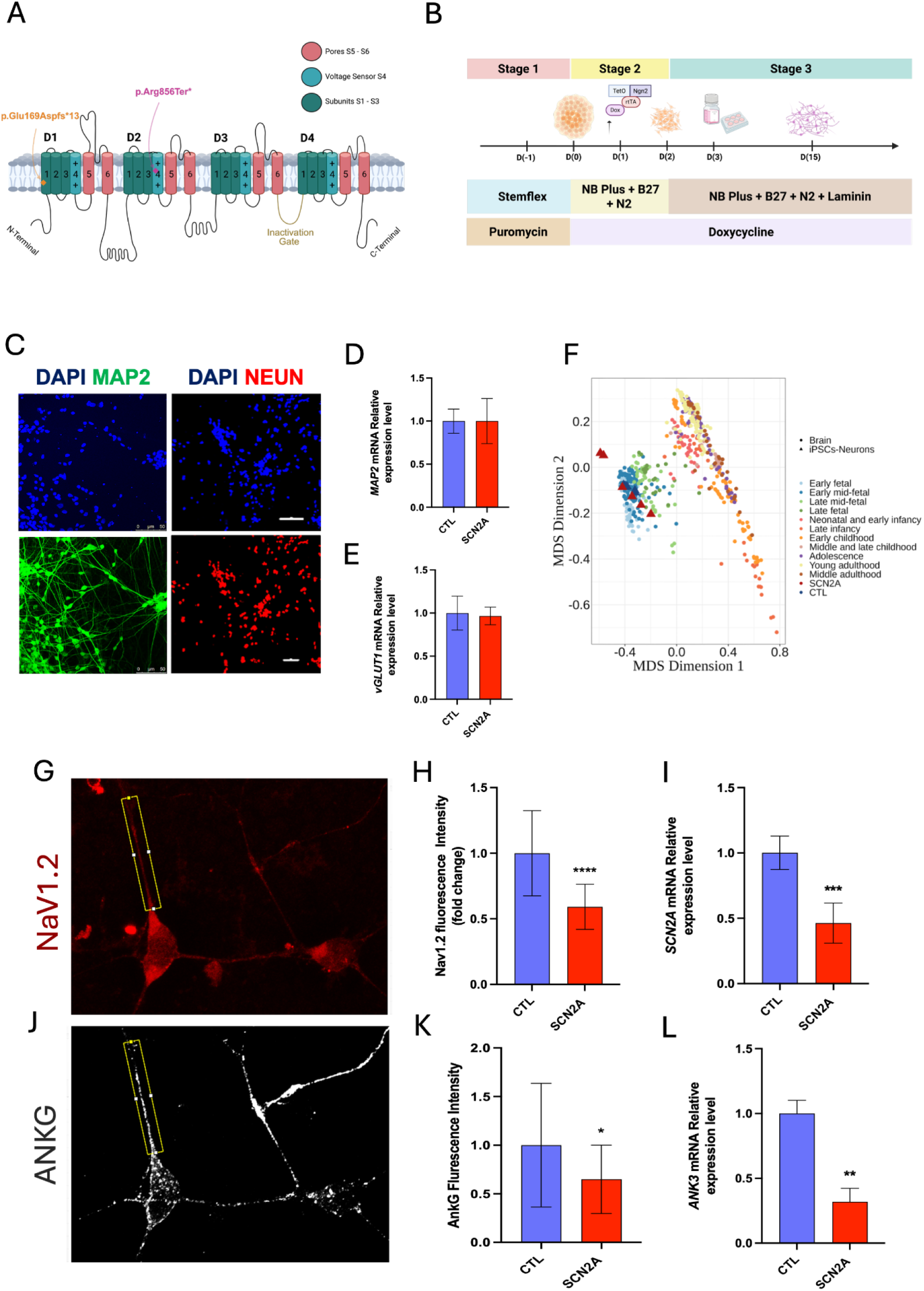
Characterization of hiPSC-derived neurons and altered *SCN2A* expression in LoF variants. **A** Schematic representation of the NaV1.2 (*SCN2A*) channel showing the positions of the patient-specific variants: the frameshift variant (c.507delA) located in Domain I and the nonsense variant (c.2566C>T) located in Domain II. **B** Overview of the NGN2-based neuronal differentiation protocol used to generate human induced neurons from hiPSCs. **C** Representative immunofluorescence images of MAP2- and NEUN-positive cells confirming neuronal identity (scale bars, 50 μm). **D–E** Relative mRNA expression of *MAP2* and *vGLUT1* showing no significant differences between control and patient neurons. **F** Multidimensional scaling (MDS) analysis showing that NGN2-induced neurons cluster with human cortical developmental transcriptomes. **G** Representative NaV1.2 immunostaining highlighting reduced axon initial segment (AIS) signal in patient neurons. **H** Quantification of NaV1.2 fluorescence intensity (****p < 0.0001). **I** Relative *SCN2A* mRNA expression showing significant reduction in *SCN2A* LoF neurons (***p < 0.001). **J** Representative Ankyrin-G (ANKG) immunostaining at the AIS. **K** Quantification of ANKG fluorescence intensity (*p < 0.05). **L** Relative *ANK3* mRNA expression (**p < 0.01).

Quantitative RT-PCR analysis revealed a significant reduction in *SCN2A* expression at transcript levels (Figure 1I, p < 0.001), consistent withYes NMD-mediated degradation of PTC-bearing transcripts. Immunostaining confirmed that NaV1.2 is enriched at the AIS (Figure 1G), but we found a significant reduction in NaV1.2 fluorescence intensity in *SCN2A* LoF neurons (Figure 1H, p < 0.0001), indicating a pronounced loss of NaV1.2 protein at the AIS. Given the critical role of NaV1.2 at the AIS, we examined Ankyrin-G (ANKG), the scaffolding protein responsible for anchoring voltage-gated sodium channels at the AIS. AnKG localization was clearly observed at the AIS (Figure 1J), and both AnKG protein fluorescence intensity (Figure 1K, p < 0.05) and ANK3 mRNA expression (Figure 1L, p < 0.01) were significantly reduced in *SCN2A* LoF neurons.

Given that *SCN2A* encodes a complex repertoire of transcript isoforms with distinct coding potential and NMD susceptibility, we next asked how *SCN2A* LoF variants alter this isoform landscape. We analyzed neurons generated from unaffected control and *SCN2A* LoF hiPSCs, differentiated under identical conditions to isolate variant-driven effects on transcript fate.

Across patient-derived *SCN2A* LoF neurons, most *SCN2A* transcript isoforms exhibited marked downregulation relative to controls, with a magnitude and pattern that differed across the two variants (Figure 2A). Isoforms predicted to be sensitive to NMD, including *SCN2A*-211 and *SCN2A*-215, were consistently among the most reduced across all patient lines, supporting an active degradation mechanism. Neurons carrying nonsense variants (SCN2A-1 and SCN2A-3) showed the strongest reductions, often exceeding −4 to −5 log2FC for key isoforms (e.g., *SCN2A*-204, *SCN2A*-206). In contrast, the frameshift variant (SCN2A-2) produced an attenuated, and more uniform decrease (approximately −0.2 to −0.8 log2FC across isoforms), suggesting differences in NMD sensitivity or transcript stability depending on the location and nature of the mutation. These patterns clarify that the pathogenic variants do not simply reduce overall abundance but remodel the isoform landscape of *SCN2A*, with selective depletion of NMD-targeted transcripts.

**Figure 2.**
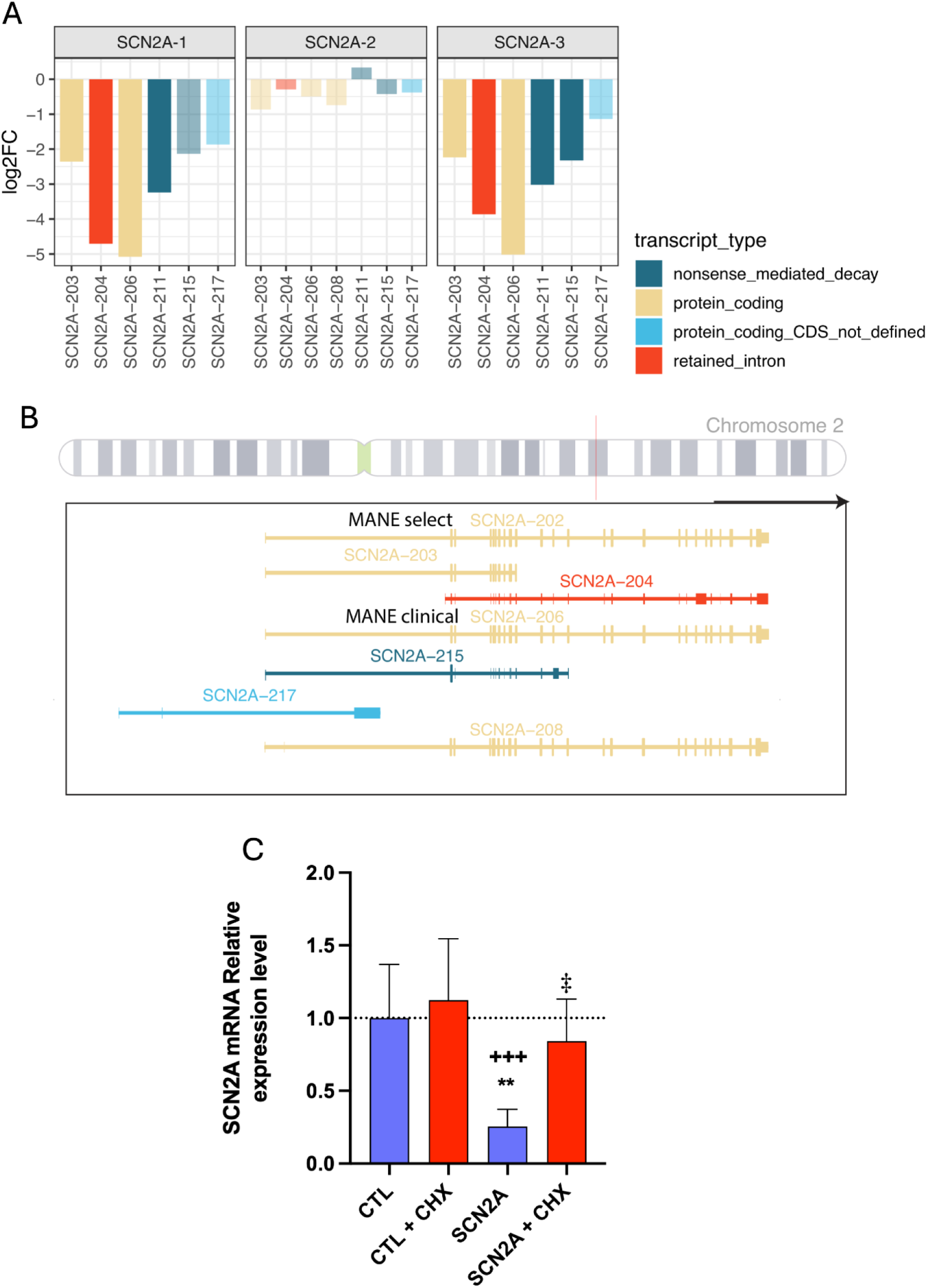
Isoform-resolved *SCN2A* expression and NMD validation. **A** Differential expression profiles of *SCN2A* transcript isoforms (*SCN2A*-203, *SCN2A*-204, *SCN2A*-206, *SCN2A*-208, *SCN2A*-211, *SCN2A*-215, *SCN2A*-217) across three patient-derived *SCN2A* LoF neurons compared with controls. Bars indicate log2 fold-change (log2FC). Isoform categories are color-coded as nonsense-mediated decay (dark teal), protein-coding (tan), protein-coding-CDS-not-defined (light blue), and retained-intron (red/orange). **B** Genomic representation of *SCN2A* isoforms, including Matched Annotation from Ensembl and ncbi (MANE)-select and MANE-clinical transcripts. **C** qPCR quantification of *SCN2A* mRNA levels under four conditions: CTL, CTL + CHX, *SCN2A*, and *SCN2A* + CHX. Cycloheximide (CHX) treatment blocks translation and reveals NMD-sensitive transcripts. *SCN2A* + CHX shows significantly increased *SCN2A* mRNA levels, relative to untreated *SCN2A* neurons, consistent with active NMD-mediated degradation.

A genomic representation of the major *SCN2A* isoforms, including Matched Annotation from Ensembl and ncbi (MANE)-select and MANE-clinical transcripts (Figure 2B), further highlights that several high-confidence protein-coding isoforms undergo substantial depletion in patient-derived neurons, reinforcing the disruption of canonical *SCN2A* transcriptional architecture.

To directly test whether NMD mediates the observed isoform reduction, we quantified *SCN2A* mRNA levels following CHX treatment, which inhibits translation and blocks NMD (Figure 2C). In control neurons, CHX treatment produced only a modest, nonsignificant increase in *SCN2A* expression, consistent with minimal basal NMD activity. In contrast, *SCN2A* LoF neurons displayed markedly reduced baseline *SCN2A* levels, and CHX treatment yielded a significant rescue. Although expression did not fully return to control levels, the magnitude of the rescue provides strong functional evidence for active NMD-mediated degradation of *SCN2A* transcripts in *SCN2A* LoF neurons. These findings indicate that a substantial fraction of *SCN2A* transcripts in the *SCN2A* LoF neurons is normally degraded by NMD, supporting an active role of this pathway in mediating *SCN2A* transcript loss.

### *SCN2A* haploinsufficiency alters AIS morphology and dendritic organization

Having established that *SCN2A* LoF produces isoform-specific downregulation and activates NMD-mediated degradation, we next examined whether these molecular defects translate into measurable alterations in neuronal structure and cellular organization. Because NaV1.2 is highly enriched at AIS and contributes to excitability and axonal identity, AIS morphology served as a logical starting point for assessing cell-level consequences of *SCN2A* haploinsufficiency.

Immunofluorescence analysis revealed AIS disorganization in patient-derived neurons (Figure 3A). *SCN2A* LoF neurons exhibited a significantly shorter AIS, together with a pronounced reduction in PanNav signal intensity (Figure 3B–C), indicating reduced sodium channel density at this compartment. These structural impairments are consistent with the molecular depletion of *SCN2A* transcripts observed in Figure 2 and support the idea that reduced NaV1.2 availability compromises AIS assembly and stability. *SCN2A* LoF neurons displayed fewer dendritic intersections across multiple radii (Figure 3D), and total arborization was significantly reduced compared with controls (Figure 3E-F).

**Figure 3.**
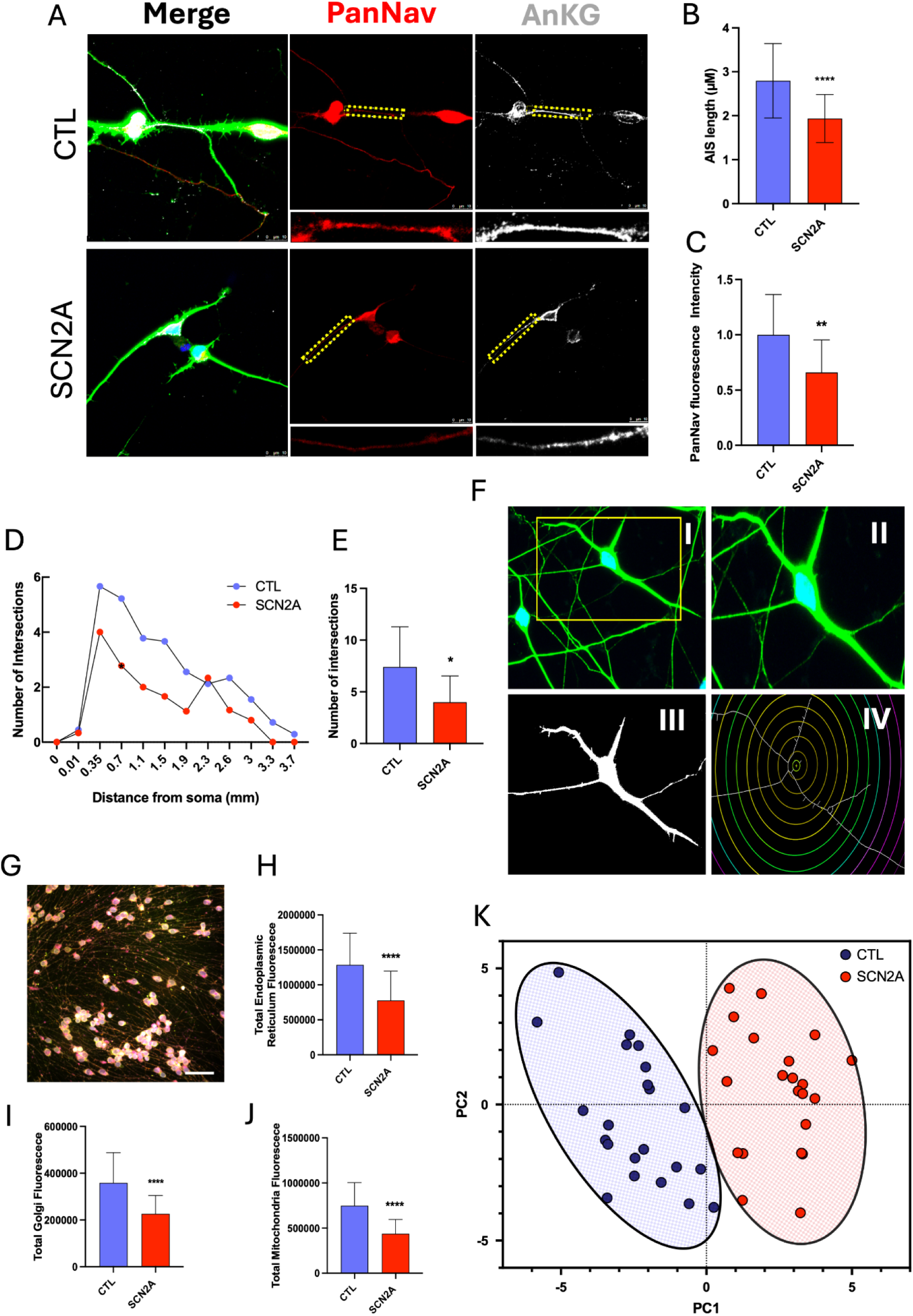
*SCN2A* haploinsufficiency disrupts AIS structure, dendritic morphology, and organelle integrity. **A** Immunofluorescence of control (CTL) and *SCN2A* LoF neurons stained for PanNav (red) and Ankyrin-G (AnKG, white) highlighting the axon initial segment (AIS; yellow dashed lines). **B** AIS length is significantly reduced in *SCN2A* LoF neurons (****P < 0.0001). **C** PanNav fluorescence intensity at the AIS is decreased (**P < 0.01). **D** Sholl profiles reveal reduced dendritic complexity in *SCN2A* LoF neurons. **E** Quantification of total intersections confirms a significant reduction (*P < 0.05). **F** Workflow for morphological extraction from fluorescence images (raw image, segmentation, and Sholl overlays). **G** Representative composite of Cell Painting channels used for high-content organelle profiling. **H–J** *SCN2A* LoF neurons exhibit markedly reduced fluorescence intensity for endoplasmic reticulum (****P < 0.0001), Golgi apparatus (****P < 0.0001), and mitochondria (****P < 0.0001). **K** PCA of Cell Painting–derived features shows clear separation between CTL and *SCN2A* LoF neurons, indicating a distinct morphological–organelle phenotype associated with *SCN2A* haploinsufficiency.

To determine whether *SCN2A* haploinsufficiency also perturbs subcellular homeostasis, we performed Cell Painting, a multiplexed high-content imaging assay that captures morphological and organelle-related features at single-cell resolution (Figure 3G). *SCN2A* LoF neurons showed reductions in endoplasmic reticulum, Golgi, and mitochondrial fluorescence intensities (Figure 3H–J), indicating altered organelle abundance or organization. These changes likely reflect broader defects in trafficking, membrane dynamics, and metabolic function. PCA of all Cell Painting variables revealed a segregation between *SCN2A* LoF and control neurons (Figure 3K), demonstrating that the combination of AIS abnormalities, dendritic simplification, and organelle perturbations forms a coherent and discriminative cellular phenotype associated with *SCN2A* haploinsufficiency.

### Transcriptomic profiling uncovers convergent dysregulation of synaptic and neurodevelopmental programs

To extend the molecular alterations observed at the structural and cellular levels, we next performed bulk RNA-seq on day-15 neurons derived from hiPSC lines. Consistent with the AIS disorganization, reduced dendritic complexity, and organelle vulnerability observed in Figures 2 and 3, transcriptome-wide analysis revealed a coherent remodeling of neuronal transcriptional programs.

First, PCA demonstrated a separation between *SCN2A* LoF and control neurons (Figure 4A), indicating that *SCN2A* LoF establishes a global transcriptional state distinct from canonical transcriptional maturation trajectory, which aligns with the morphological and subcellular phenotypes described previously. Differential expression analysis identified a focused set of dysregulated genes (Figure 4B), many of which are directly involved in synaptic architecture, neuronal excitability, and neurodevelopmental signaling. Several top hits mapped to ion channel regulation, glutamatergic synapse components, and axonal specification modules. GO Biological Process enrichment revealed downregulation of pathways associated with synaptic signaling, trans-synaptic communication, neuron projection development, and regulation of membrane potential (Figure 4C). Human Phenotype Ontology enrichment further highlighted phenotypic categories that mirror the clinical spectrum associated with *SCN2A*-related NDDs, including seizure susceptibility, abnormal social behavior, and neurodevelopmental delay (Figure 4D).

Consistent with these functional annotations, GO Cellular Component terms showed strong enrichment of postsynaptic and synaptic membrane compartments, neuronal projections, axonal structures, and ion channel complexes (Figure 4E). GO Molecular Function categories pointed to altered activity of voltage-gated sodium, potassium and calcium channels, as well as neurotransmitter receptor signaling (Figure 4F), reinforcing the ion-channel-centered architecture of *SCN2A* biology.

Finally, hierarchical clustering of the top DEGs produced a coherent transcriptional signature that discriminated *SCN2A* LoF neurons from control neurons (Figure 4G). Gene modules associated with excitatory neurotransmission, axonal maturation, and synaptic alignment were consistently reduced. Together, these results show that *SCN2A* haploinsufficiency interferes not only with AIS organization and neuronal morphology but also with the transcriptional mechanisms that support synaptic connectivity and neuronal excitability.

### Isoform-resolved transcriptomics uncovers NMD engagement and lncRNA remodeling

To further investigate the regulatory layers underlying the transcriptional disruptions identified in Figure 4, we next performed RNA-seq at the transcript level using samples with 45M read-depth paired-end libraries, focusing on isoform usage, NMD activity, and lncRNA-mediated regulatory networks. This approach enabled quantification of differentially expressed transcripts (DETs) across all major RNA biotypes; including protein-coding isoforms, retained-intron transcripts, NMD-targeted isoforms, and long non-coding RNAs (lncRNAs), providing a higher-resolution view of transcriptional remodeling in *SCN2A* LoF neurons.

**Figure 4.**
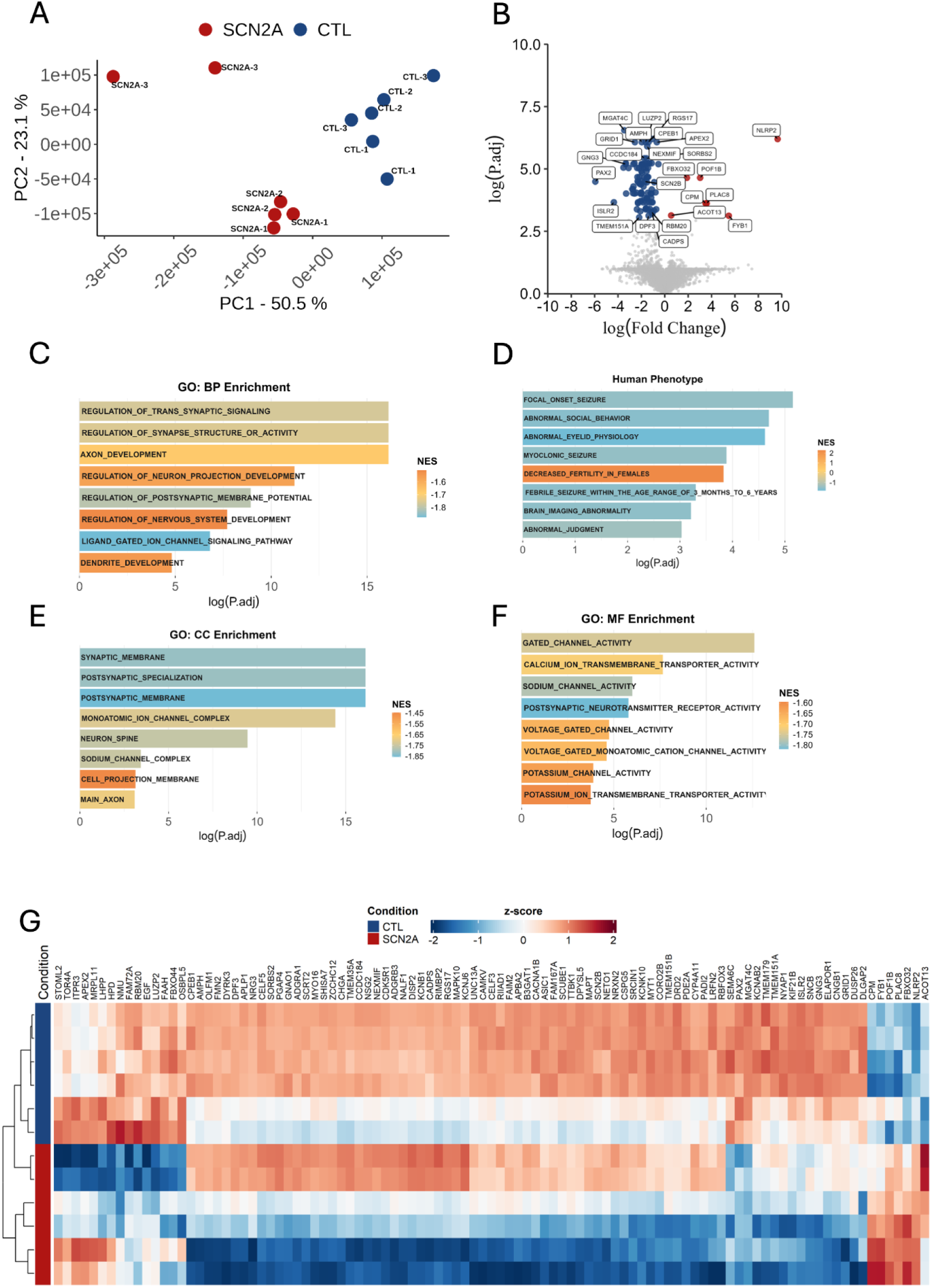
Global transcriptomic profiling of *SCN2A*-haploinsufficient neurons reveals coordinated disruption of synaptic, ion channel, and neurodevelopmental programs. **A** PCA of RNA-seq datasets showing a segregation between control (CTL) and *SCN2A* LoF patient neurons along PC1, indicating a genotype-driven shift in global transcriptional signatures. **B** Volcano plot highlighting DEGs between CTL and *SCN2A* LoF groups (FDR < 0.05; |log2FC| > 1). DEGs related to neuronal excitability and synaptic organization are indicated. **C** Gene Ontology—Biological Process (GO-BP) enrichment analysis showing significant downregulation of pathways related to synaptic signaling, trans-synaptic communication, neuron projection development, and regulation of membrane potential (FDR < 0.05). **D** Human Phenotype Ontology (HPO) enrichment demonstrating overrepresentation of seizure-related, behavioral, and neurodevelopmental phenotypes associated with *SCN2A* dysfunction (FDR < 0.05). **E** GO—Cellular Component (GO-CC) enrichment revealing significant enrichment of postsynaptic density, synaptic membrane, ion channel complexes, neuronal projections, and dendritic structures (FDR < 0.05). **F** GO—Molecular Function (GO-MF) enrichment highlighting altered activity of voltage-gated sodium, potassium, and calcium channels, as well as neurotransmitter receptor binding and signaling (FDR < 0.05). **G** Heatmap of the top dysregulated genes (hierarchical clustering using Pearson correlation) illustrating a coherent transcriptional signature in *SCN2A* LoF neurons characterized by coordinated downregulation of synaptic, ion channel, and neurodevelopmental gene modules. All statistical analyses were performed using DESeq2 with Benjamini–Hochberg correction.

**Figure 5.**
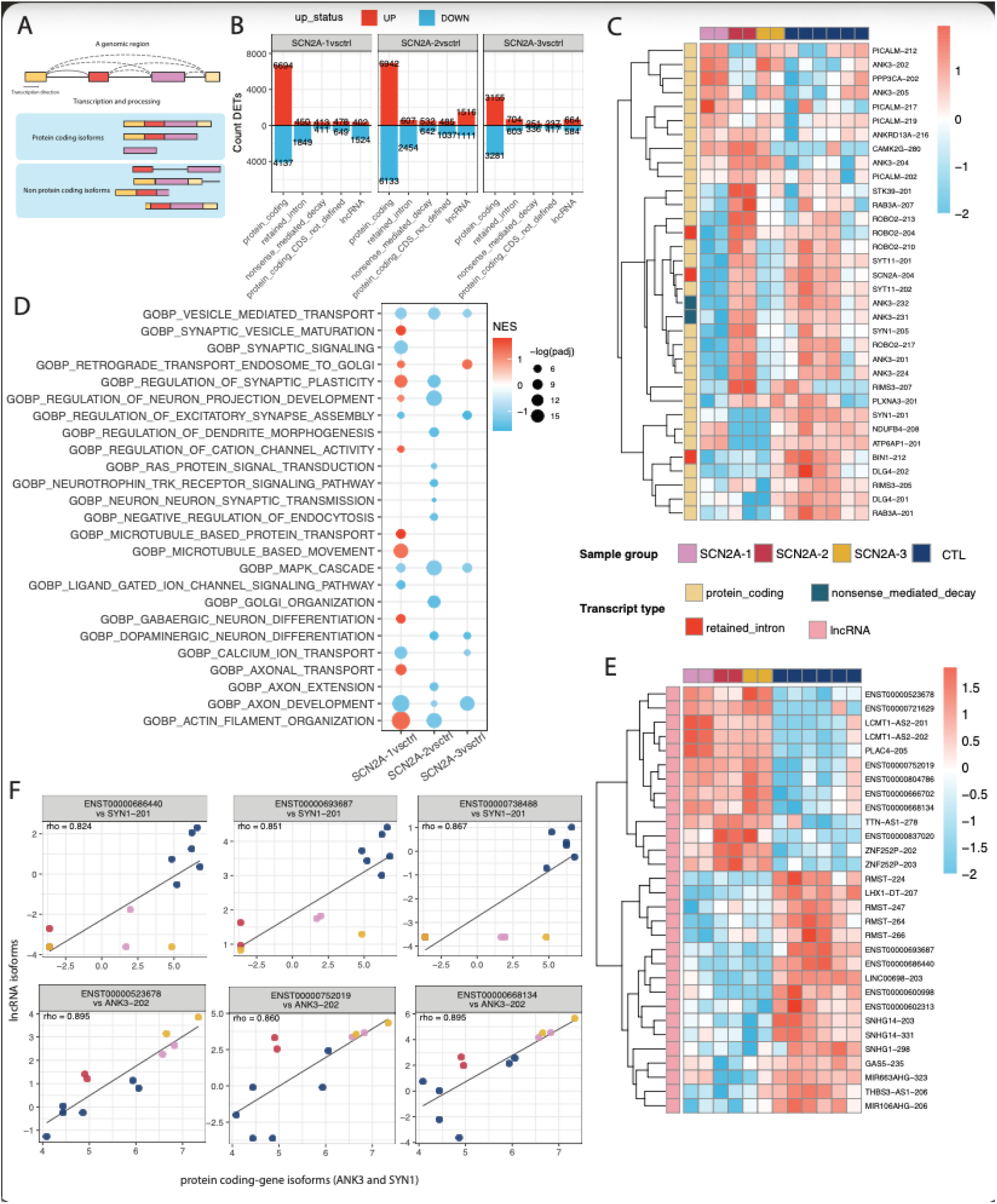
Transcript-level remodeling, NMD activation and lncRNA–protein-coding correlations in *SCN2A*-haploinsufficient neurons. (**A)** Schematic representation of transcript biotypes arising from a single genomic locus, illustrating alternative splicing generating protein-coding, retained-intron, NMD-targeted and non-coding isoforms. (**B)** Bar plots showing the number of differentially expressed transcripts (DETs) in each *SCN2A* LoF patient line (SCN2A-1, SCN2A-2, SCN2A-3) relative to controls. DETs (|log₂FC| > 1; adj. *p* < 0.05) are grouped by transcript biotype and direction of regulation (up/down). **(C)** Heatmap of selected DET protein-coding isoforms, showing scaled logCPM expression across control and *SCN2A* LoF lines. Transcript type and sample group are indicated in the side and top annotations. **(D)** Gene Ontology Biological Process enrichment (protein-coding transcripts only). Dot size represents gene count and color indicates normalized enrichment score (NES), revealing consistent enrichment for synaptic signaling, vesicle trafficking, axon development, cytoskeletal organization and calcium channel–related processes. **(E)** Heatmap of differentially expressed lncRNA isoforms, highlighting distinct expression profiles across *SCN2A* LoF lines compared to controls. Biotype and group annotations correspond to panel C. **(F)** Representative correlations between lncRNA isoforms and their paired protein-coding transcripts (Spearman ρ > 0.8, *p* < 0.01). Each point represents one sample (CTL, SCN2A-1, SCN2A-2, SCN2A-3); solid lines show linear fits. Examples include lncRNAs correlated with SYN1-201 and ANK3 protein-coding isoforms. Full correlation sets are provided in Supplementary Figure XX and Supplementary Table Sx.

To dissect transcriptional remodeling beyond gene-level changes, we examined DETs across annotated isoform biotypes. Unlike the predominantly downregulated profile observed at the gene level (Figure 4), isoform-resolved analysis revealed a more heterogeneous landscape, with both upregulated and downregulated DETs across protein-coding transcripts, NMD-sensitive isoforms, retained-intron isoforms, and lncRNAs (Figure 6A–C). Importantly, the net proportion of upregulated DETs varied by biotype, indicating that isoform remodeling in *SCN2A* LoF neurons does not follow a uniform directionality.

Protein-coding transcripts showed a modest predominance of upregulated isoforms, whereas retained-intron and predicted NMD-targeted isoforms exhibited a mixed response, with subsets of transcripts increased and others decreased in abundance. This pattern is consistent with competing post-transcriptional processes, including increased degradation of certain PTC-containing isoforms while others may accumulate due to altered splicing or reduced translation-coupled decay. Thus, our data suggests that *SCN2A* haploinsufficiency induces biotype-specific and isoform-specific regulation, rather than global downregulation of any single class. All three *SCN2A*-haploinsufficient lines exhibited a tendency toward upregulated transcripts, including an increased representation of NMD-associated transcripts. Notably, the two lines carrying the same nonsense variant (SCN2A-1 and SCN2A-3; p.Arg856Ter) exhibited highly similar DET profiles across biotypes, whereas the frameshift line (SCN2A-2; p.Glu169Aspfs13) showed a more attenuated and compositionally distinct pattern (Figure 6B).

Heatmap analyses further illustrated distinct expression patterns across transcript types (Figure 6C–E). Protein-coding DETs segregated *SCN2A* and control samples into clearly separated clusters, while retained-intron and NMD-related transcripts exhibited coherent downregulation consistent with increased RNA degradation pressure. Enrichment analysis performed on protein-coding DETs (Figure 6D) revealed strong convergence on neuronal and synaptic pathways, including synapse organization, vesicle trafficking, calcium handling, neuron projection morphogenesis, and axon guidance.

A notable finding was that lncRNA DETs were strongly enriched for upregulated transcripts, suggesting broad reorganization of non-coding RNA programs. Many of these lncRNA isoforms have no prior characterization and have not been associated with any biological condition, suggesting the emergence of previously unrecognized regulatory signatures in *SCN2A* LoF neurons. As lncRNAs undergo splicing and often display low conservation at splice junctions, they typically generate a larger number of transcript isoforms compared to protein-coding genes (Deveson et al., 2018). They are known to influence the expression of other RNAs in their vicinity but also in completely distinct chromosomal territories (Mattick et al., 2023a). To explore potential regulatory relationships, we performed correlation analyses between lncRNA expression and protein-coding isoforms. Across lines, we identified multiple high-confidence correlation pairs (|r| > 0.8, FDR < 0.05) involving neurodevelopmental and synaptic genes (Figure 6D–F). Notably, SYN1 was strongly correlated with previously unnamed lncRNA isoforms (Figure 6F; Supplementary Figure S4), suggesting that lncRNA-mediated mechanisms may contribute to its enhanced expression in *SCN2A* LoF neurons.

We also observed extensive isoform remodeling within the ANK3 locus. ANK3 encodes a large set of protein-coding isoforms with diverse neuronal functions. Among these, ANK3-202 and ANK3-204 emerged as consistently upregulated across all *SCN2A* LoF neurons (Figure 6C; Supplementary Figure S3), and both isoforms are shorter than the canonical MANE-select isoform ANK3-201. Furthermore, ANK3-202 displayed the highest number of correlated lncRNA isoforms, including both named and unnamed transcripts (Figure 6F; Supplementary Figure S4), reinforcing the possibility that regulatory interactions between lncRNAs and specific protein-coding isoforms shape the transcriptomic landscape of *SCN2A* LoF neurons. These findings demonstrate that *SCN2A* LoF induces widespread transcriptomic remodeling extending beyond protein-coding genes to encompass alternative splicing, NMD-targeted isoforms, and non-coding RNA networks.

## Discussion

*SCN2A* is one of the most reproducible monogenic contributors to NDDs, with LoF variants typically presenting as ASD and global developmental delay without seizures (Sanders et al., 2015, 2018; Wolff; Brunklaus; Zuberi, 2019). Its dosage is particularly critical during early corticogenesis, when NaV1.2 supports axonal polarity, early excitability and the establishment of nascent excitatory circuits (Spratt et al., 2019). NGN2-induced human glutamatergic neurons adopt a fetal-like transcriptional identity (Lin et al., 2021), making them a relevant model to study *SCN2A*-NDDs. Here, by using three patient-derived hiPSCs harboring E169Dfs*13 (SCN2A-2) or R856X (SCN2A-1 and SCN2A-3) we identified a convergent cellular phenotype characterized by isoform-specific reductions consistent with NMD activity, AIS shortening and PanNav loss, coordinated remodeling of synaptic and axonal pathways, and widespread alterations in lncRNA and transcript isoform usage. These findings suggest that the consequences of NaV1.2 loss extend beyond ion-channel dysfunction to encompass multilayered disruptions in neuronal architecture and gene regulatory networks. Importantly, the phenotypic convergence across three genetically distinct patient lines, argues against line-specific artifacts and supports *SCN2A* as the primary driver of the observed alterations.

Supporting NMD as the primary mechanism driving haploinsufficiency, all three patient-derived lines converged on reductions of *SCN2A* mRNA that were partially rescued by CHX treatment (Asadollahi et al., 2023; Jaffrey; Wilkinson, 2018; Le Hir et al., 2001; Nagy; Maquat, 1998). Isoform-resolved analyses further clarified how this emerges at the transcript level. Across all patient-derived lines, NMD-sensitive isoforms - particularly *SCN2A*-211 and *SCN2A*-215 - showed the most pronounced reductions, fully consistent with their annotation as NMD-targeted transcripts in Ensembl/GENCODE (Frankish et al., 2021). This pattern mirrors canonical NMD responses reported for truncating *SCN2A* variants (Asadollahi et al., 2023) and aligns with studies showing reduced NaV1.2 expression and impaired firing in *SCN2A* LoF states (Satterstrom et al., 2020). Although earlier studies reported deficits in excitability and network activity in *SCN2A* LoF neurons (Brown et al., 2023; Tamura et al., 2025), our results extend this literature by resolving isoform-level mechanisms that clarify how haploinsufficiency emerges and propagates across regulatory and structural layers.

Importantly, degradation efficiency varied between patient lines, with NMD acting more strongly in SCN2A-1 and SCN2A-3 than in SCN2A-2, suggesting that exon–EJC geometry and transcript configuration may modulate haploinsufficiency severity. This position-dependent variation aligns with recent findings from Al Saneh and colleagues (2025), who demonstrated in mouse models that an early coding sequence PTC (Y84X) engages partial NMD, whereas a terminal-exon PTC (R1627X) escapes decay entirely - illustrating that PTC position is a critical determinant of transcript fate (Saneh et al., 2025). Prior work has shown that NMD efficiency is shaped by PTC position, exon–junction complex geometry and transcript architecture (Ogiwara et al., 2018).

Downstream of these transcript-level disruptions, the AIS emerged as a point of phenotypic convergence. All patient-derived lines exhibited reductions in NaV1.2, pan-Nav expression, Ankyrin-G and *ANK3* mRNA, accompanied by AIS shortening. Because AIS assembly depends on reciprocal stabilization between NaV1.2 and Ankyrin-G (Kole; Stuart, 2012; Lorincz; Nusser, 2008), *SCN2A* dosage loss may compromise this scaffold, potentially affecting axonal polarity, excitability and downstream synaptic maturation. The structural alterations we observed, reduced NaV1.2 density, shortened AIS, and decreased Ankyrin-G, represent well-established morphological predictors of altered excitability, as demonstrated in multiple studies of *SCN2A* LoF models (Asadollahi et al., 2023; Chen et al., 2024; Spratt et al., 2019; Tamura et al., 2025).

Structural alterations, however, extended beyond the AIS. Reduced dendritic arborization, diminished ER/Golgi/mitochondrial signatures and robust PCA separation of patient-derived neurons from controls indicate broader disturbances in intracellular organization and trafficking, features previously associated with altered neuronal development in ASD (Marchetto et al., 2017; Rossignol; Frye, 2012). Perturbations at cytoskeletal–organelle interfaces are known to reshape synapse composition (Dent; Gupton; Gertler, 2011; Reshetniak et al., 2025), and these alterations may represent either direct consequences or secondary adaptations to NaV1.2 loss. Transcriptomic enrichment for synaptic signaling, ion-channel function, morphogenesis and neurotransmitter pathways aligned well with these structural findings and mirrored developmental convergence observed in ASD risk genes during early corticogenesis (Cabana-Domínguez et al., 2023; Paranjapye et al., 2025; Satterstrom et al., 2020).

A further dimension of our dataset involves widespread alterations in noncoding RNA classes. Extensive changes in NMD targets and previously unannotated lncRNA isoforms indicate that *SCN2A* haploinsufficiency perturbs not only protein-coding transcripts but also broader RNA landscapes. Neurodevelopmentally relevant lncRNAs display low splicing-site conservation and high structural diversity (Fico et al., 2019; Mattick et al., 2023b), and have been implicated in chromatin organization and post-transcriptional control in other contexts (Statello et al., 2021). We observed correlations between specific lncRNA isoforms, ANK3 variants, and synaptic genes such as SYN1. These associations raise the possibility that lncRNA alterations may be part of the broader transcriptional response to NaV1.2 loss, consistent with ASD multi-omic studies demonstrating concurrent changes in splicing and lncRNA expression (Parikshak et al., 2016). While the correlative nature of these associations precludes causal inference, the consistent co-variation between specific lncRNA isoforms and neurodevelopmentally relevant genes, including SYN1 and ANK3, positions these non-coding transcripts as candidate regulators warranting functional investigation. Future studies employing targeted lncRNA perturbation will be required to establish mechanistic roles.

## Limitations

Our model is constrained by the accelerated developmental trajectory of NGN2-induced neurons, which recapitulate early corticogenesis but not later phases of circuit refinement. The use of non-isogenic controls introduces background genetic variability, although phenotypic convergence across patient lines partially mitigates this concern. Bulk RNA-seq limits definitive isoform quantification. Electrophysiological recordings were not performed, leaving functional correlates grounded in established literature rather than direct measurement. While direct functional readouts would complement our findings, we note that: (1) functional deficits in *SCN2A* LoF neurons are well-established in the literature (Brown et al., 2023; Chen et al., 2025; Olivero-Acosta et al., 2025; Que et al., 2021; Spratt et al., 2019; Tamura et al., 2025; Zhang et al., 2021); (2) our structural and molecular findings align with these established functional phenotypes; and (3) the primary contribution of this work lies in defining the multi-omic molecular architecture underlying *SCN2A* haploinsufficiency rather than recapitulating functional readouts. Future studies integrating electrophysiology with multi-omic profiling would provide additional mechanistic depth. Our analyses were conducted at a single timepoint (day 15), precluding conclusions about temporal dynamics. Finally, correlations between lncRNA and protein-coding transcripts are associative, not causal, and will require targeted perturbation experiments to establish mechanistic roles. While these associations do not establish causality, they point to coordinated transcriptional modules potentially influenced by lncRNA activity, consistent with emerging evidence that lncRNAs shape post-transcriptional and splicing dynamics during neuronal maturation.

## Concluding Remarks

Our multi-omic analysis reveals that *SCN2A* haploinsufficiency perturbs neuronal development through a coordinated cascade of molecular and structural alterations. *SCN2A* LoF activates NMD and reshapes *SCN2A* isoform usage, leading to AIS shortening, reduced dendritic complexity, organelle vulnerability.. These findings refine the mechanistic landscape underlying previously described electrophysiological deficits and demonstrate that *SCN2A* LoF influences neuronal identity far beyond ion-channel dysfunction. Transcript-resolved RNA-seq adds an additional layer of insight, uncovering lncRNA signatures pointing to post-transcriptional regulation as a contributing mechanism. Together, these convergent alterations define a multilayered phenotype spanning gene dosage, isoform regulation and neuronal architecture. The delineation on how these pathways interact point to possible therapeutic directions, particularly isoform-level *SCN2A* restoration and modulation of AIS-associated regulatory networks, that merit exploration in future mechanistic and translational studies.

## Acknowledgements

We thank all members of the laboratory for their technical support, constructive discussions, and critical feedback throughout the development of this study. We also acknowledge the institutional core facilities for providing access to imaging, sequencing, and computational resources essential for data acquisition and analysis. The contribution of collaborators involved in patient recruitment, cell line generation, and methodological guidance is gratefully recognized. This work was supported by graduate fellowships and institutional funding mechanisms that enabled the execution of the experimental and bioinformatic components of the project. Part of the computational analysis was performed at the GLORIOSOS HPC cluster at the Laboratory of Genetics Biochemistry at UFMG, which is managed by Lúcio R. Queiroz, Herón Hilário and Izabela Mamede.

## Data availability statement

The entire raw RNA-seq data for the project is available at Sequence read Archive (SRA) under accession number: PRJNA1365646.

