## Supplementary Information for "Integrated multiomic profiling of *SCN2A* loss-of-function reveals widespread molecular remodeling in patient hiPSC-derived neurons"

### Supplementary Figures

**Supplementary Figure S1. Characterization and Quality Control of Patient-Derived and Control iPSC Lines**

**Supplementary Figure S2. Organelle-level morphological profiling reveals widespread structural alterations in SCN2A haploinsufficient neurons.**

**Supplementary Figure 3 ANK3 isoform-level differential expression and genomic organization.**

**Supplementary Figure 4. Correlation between ANK3 or SYN1 protein-coding isoforms and selected lncRNA isoforms.**

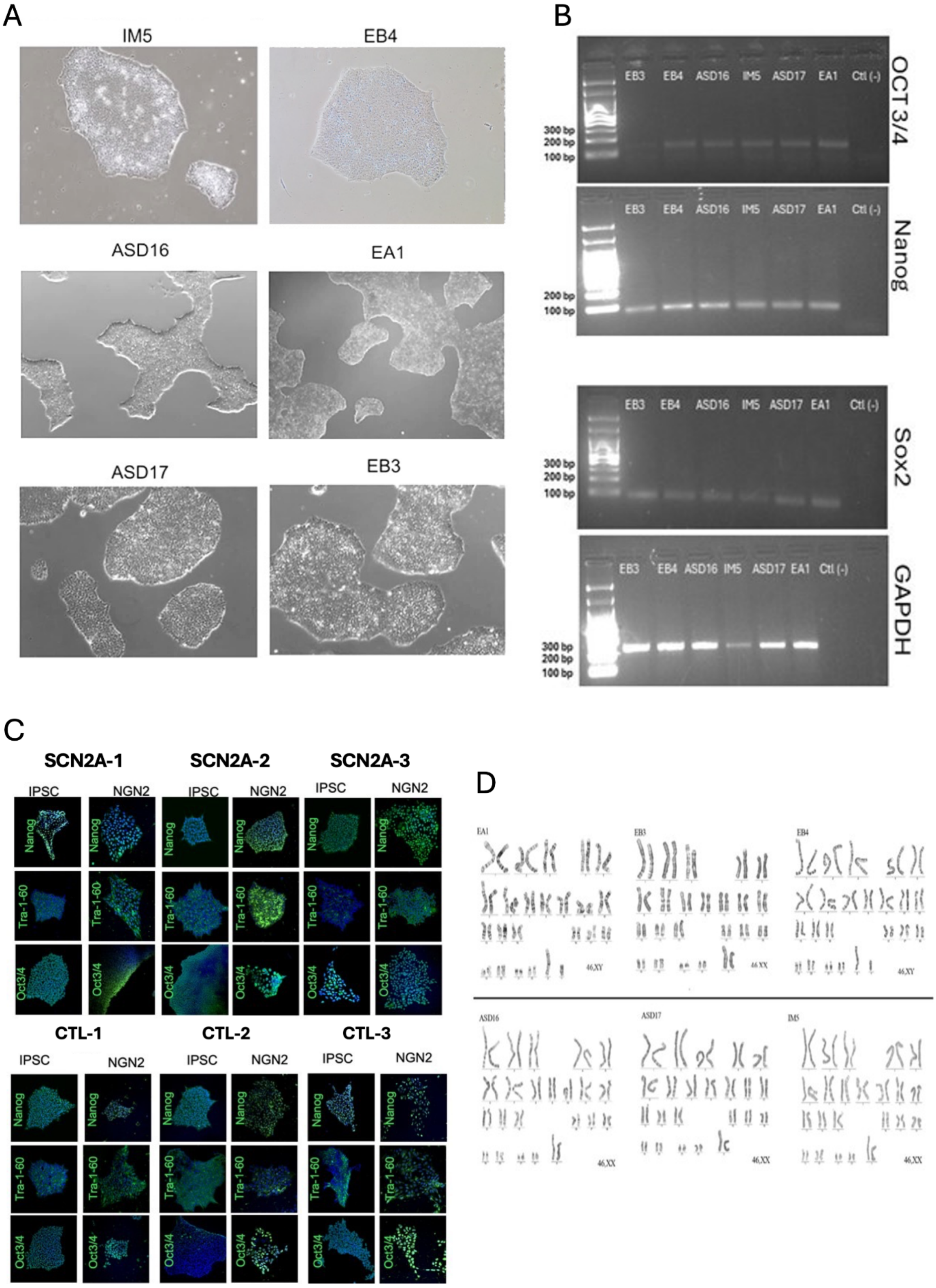


**Supplementary Figure S1. Characterization and Quality Control of Patient-**

**Derived and Control iPSC Lines (A)** Bright-field images of undifferentiated iPSC colonies. Original sample identifiers correspond to the standardized nomenclature used throughout the manuscript as follows: ASD16 (SCN2A-1), ASD17 (SCN2A-2), IM5 (SCN2A-3), EA1 (CTL-1), EB3 (CTL-2), and EB4 (CTL-3). All lines exhibit characteristic pluripotent morphology with compact colonies and high nuclear-to-cytoplasmic ratio. **(B)** RT–PCR analysis confirming expression of core pluripotency markers OCT3/4, Nanog, and Sox2 across all iPSC lines (SCN2A-1/2/3 and CTL-1/2/3). GAPDH served as an internal control. No amplification was detected in negative controls. **(C)** Immunofluorescence staining of undifferentiated iPSCs and NGN2-induced neurons (day 4) for Nanog, TRA-1-60, TRA-1-81, and SSEA4. Pluripotency markers are strongly expressed in iPSCs and appropriately downregulated after NGN2 induction, confirming robust lineage specification across all six lines. **(D)** G-banding karyotyping of the SCN2A and control iPSC lines listed above. All lines display normal chromosomal complements (46,XX or 46,XY), with no detectable structural abnormalities, supporting their suitability for downstream differentiation and functional assays.

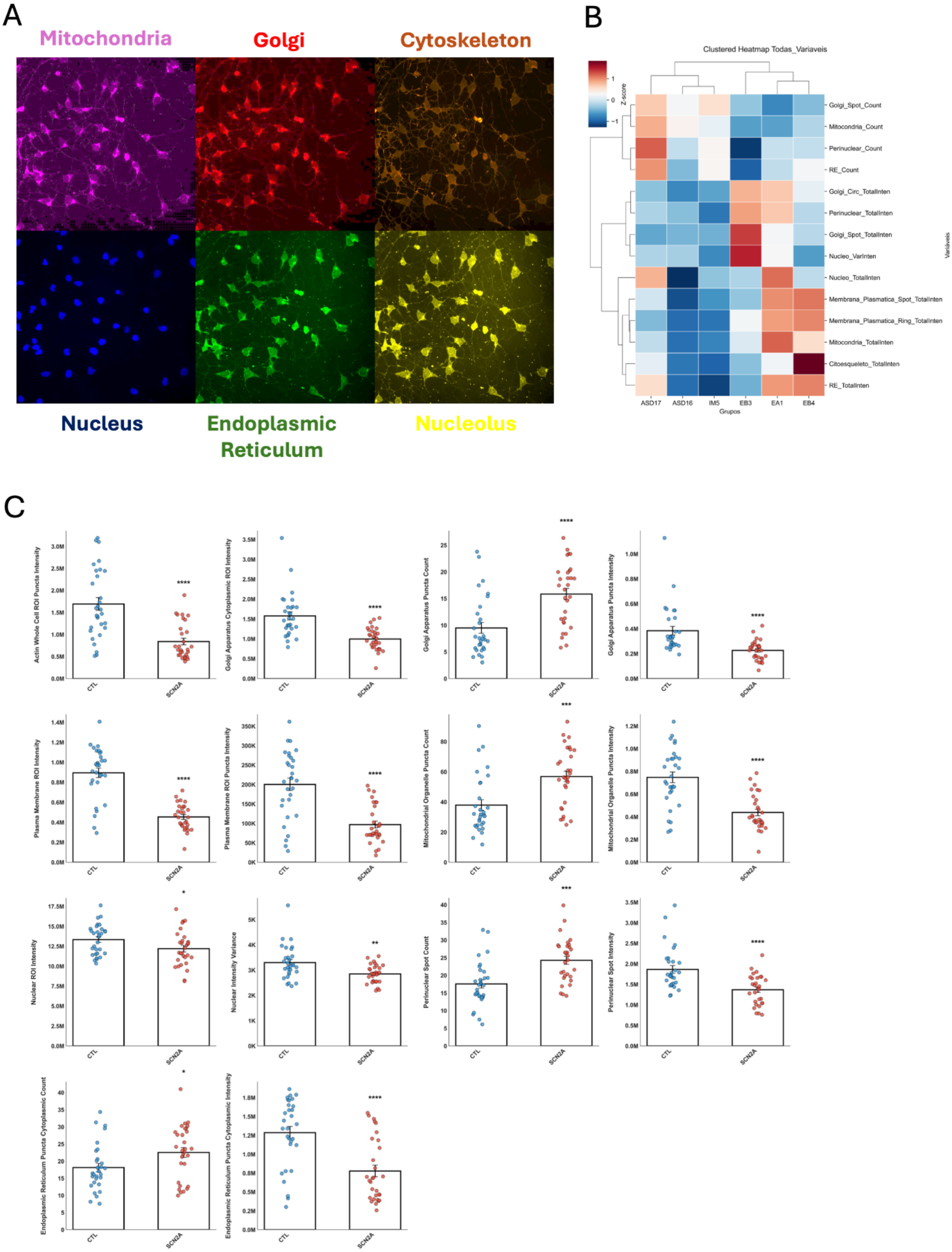
A
Mitochondria
Golgi
Cytoskeleton
Nucleus
Endoplasmic Reticulum
Nucleolus
B
Clustered Heatmap Todas_Variaveis
Z-score
Golgi_Spot_Count
Mitocondria_Count
Perinuclear_Count
RE_Count
Golgi_Circ_TotalInten
Perinuclear_TotalInten
Golgi_Spot_TotalInten
Nucleo_VarInten
Nucleo_TotalInten
Membrana_Plasmatica_Spot_TotalInten
Membrana_Plasmatica_Ring_TotalInten
Mitocondria_TotalInten
Citoesqueleto_TotalInten
RE_TotalInten
ASD17
ASD16
IM5
EB3
EA1
EB4
Grupos
Variaveis
C
Actin Whole Cell ROI Puncta Intensity
Golgi Apparatus Cytoplasmic ROI Intensity
Golgi Apparatus Puncta Count
Golgi Apparatus Puncta Intensity
Plasma Membrane ROI Intensity
Plasma Membrane ROI Puncta Intensity
Mitochondrial Organelle Puncta Count
Mitochondrial Organelle Puncta Intensity
Nuclear ROI Intensity
Nuclear Intensity Variance
Perinuclear Spot Count
Perinuclear Spot Intensity
Endoplasmic Reticulum Puncta Cytoplasmic Count
Endoplasmic Reticulum Puncta Cytoplasmic Intensity
CTL
SCN2A

**Supplementary Figure S2. Organelle-level morphological profiling reveals widespread structural alterations in SCN2A haploinsufficient neurons.(A)** Representative multiplexed Cell Painting images of NGN2-induced neurons stained for major organelles, including mitochondria (magenta), Golgi apparatus (red), cytoskeleton (orange), nucleus (blue), endoplasmic reticulum (green), and nucleolus (yellow). SCN2A loss-of-function neurons display altered cellular organization and subcellular architecture across multiple compartments.**(B)** Clustered heatmap of organelle-derived features across control and SCN2A mutant lines (CTL-1/2/3; SCN2A-1/2/3). Z-scored metrics reveal coordinated shifts in organelle number, texture, and spatial distribution, with SCN2A haploinsufficient neurons showing consistent deviations in mitochondrial organization, Golgi complexity, nucleolar features, and ER structure. **(C)** Quantification of organelle-associated features demonstrating significant differences between control and SCN2A haploinsufficient neurons. Metrics include intensity-based, texture-based, and count-based descriptors for mitochondria, the Golgi apparatus, the nucleus, the nucleolus, the cytoskeleton, and the endoplasmic reticulum. Scatter plots represent individual cells pooled across biological replicates, with mean ± s.e.m. Statistical significance was determined using unpaired two-tailed tests; *P*-values are indicated on each panel.

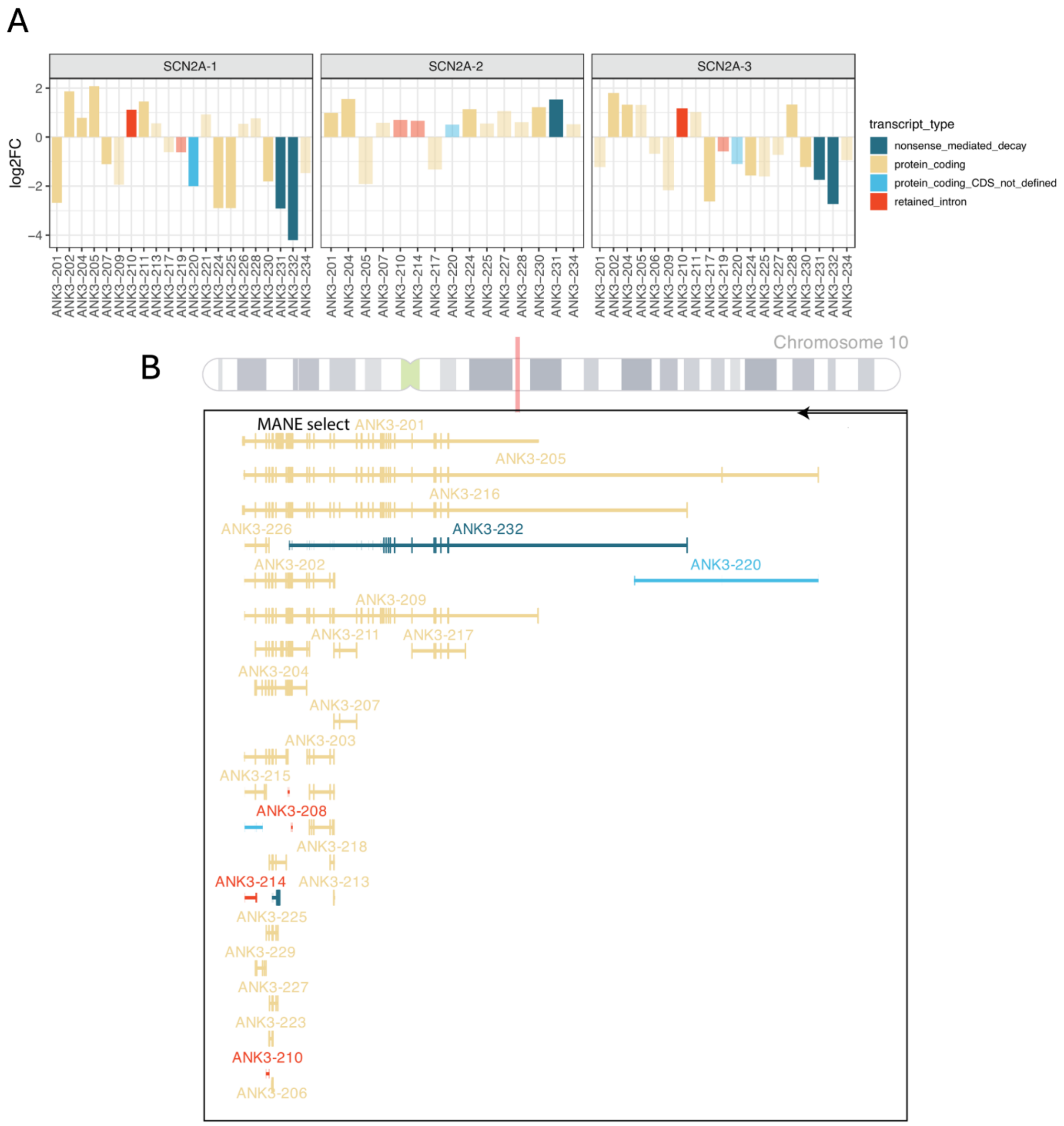


**Supplementary Figure 3 ANK3 isoform-level differential expression and genomic organization.** Bar plots (top) show log2 fold change (log2FC) for individual ANK3 transcripts in SCN2A patient-derived lines (SCN2A-1, SCN2A-2, SCN2A-3) relative to controls. Bars are colored by transcript biotype. The genomic view (bottom) displays the ANK3 locus on chromosome 10, with the MANE-select isoform ANK3-201 and additional protein-coding isoforms, highlighting the shorter upregulated isoforms ANK3-202 and ANK3-204 relative to the canonical ANK3-201.

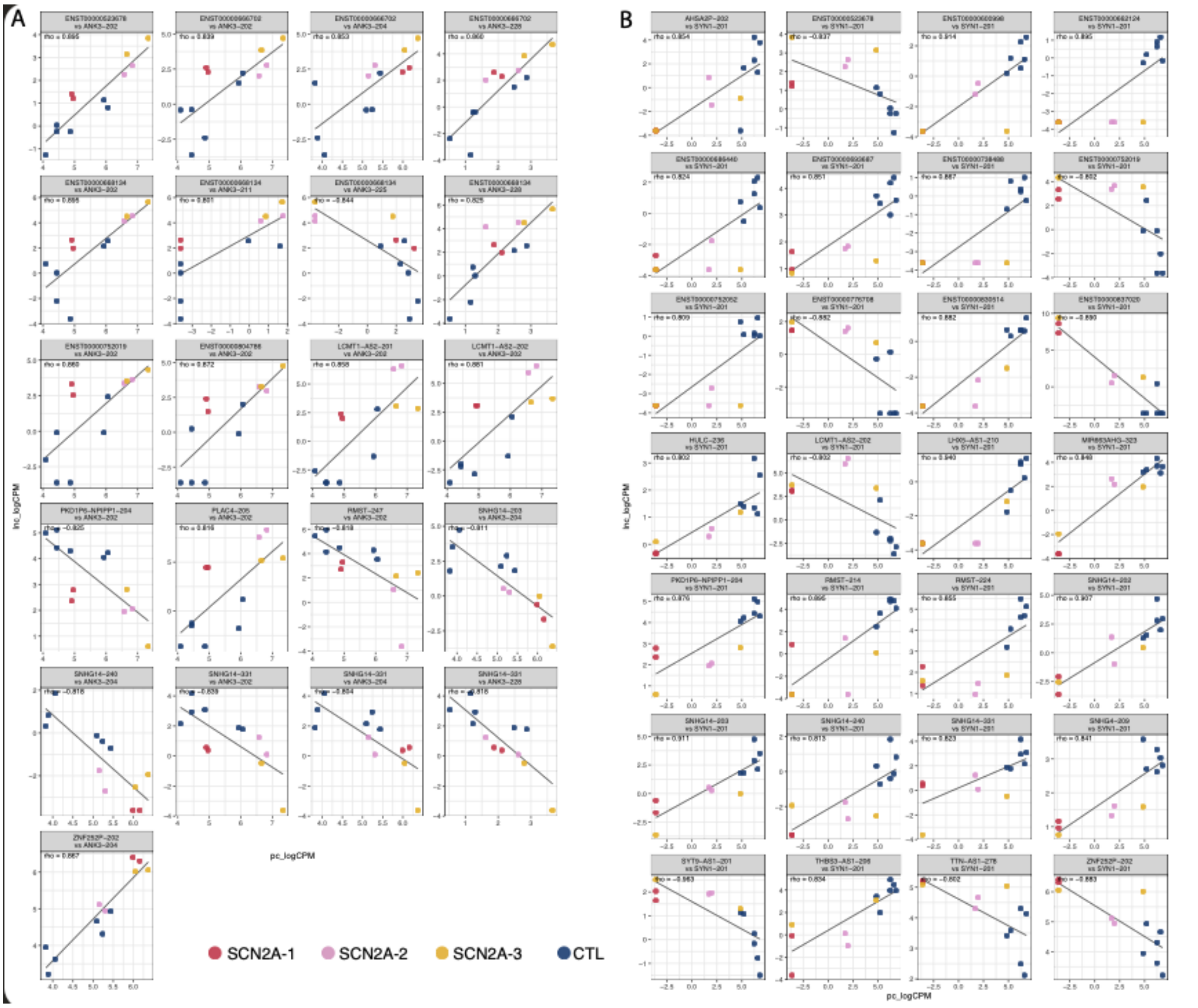


**Supplementary Figure 4. Correlation between ANK3 or SYN1 protein-coding isoforms and selected lncRNA isoforms.** (A) Scatterplots of log2 CPM expression of ANK3 protein-coding isoforms (x-axis) versus correlated lncRNA isoforms (y-axis) across control and SCN2A mutant lines. (B) Equivalent analysis for the synaptic gene SYN1 (SYN1-201) and its correlated lncRNA isoforms. Each point represents one sample (color-coded as SCN2A-1, SCN2A-2, SCN2A-3, or CTRL), solid lines show linear fits, and panel headers report the Spearman correlation coefficient (ρ) for each lncRNA–protein-coding isoform pair.

**Supplementary Table S1: DEGs table**

Supplementary Table S2: Differentially present proteins.

Supplementary Table S3: Differentially expressed Transcripts
